# Active vision in binocular depth estimation: a top-down perspective

**DOI:** 10.1101/2023.07.15.549132

**Authors:** Matteo Priorelli, Giovanni Pezzulo, Ivilin Peev Stoianov

## Abstract

Depth estimation is an ill-posed problem: objects of different shapes or dimensions, even if at different distances, may project to the same image on the retina. Our brain uses several cues for depth estimation, including monocular cues such as motion parallax and binocular cues like diplopia. However, it is still unclear how the computations required for depth estimation are implemented in biologically plausible ways. State-of-the-art approaches to depth estimation based on deep neural networks implicitly describe the brain as a hierarchical feature detector. Instead, we propose an alternative approach that casts depth estimation as a problem of active inference. We show that depth can be inferred by inverting a hierarchical generative model that simultaneously predicts the eyes projections from a 2D belief over an object. Model inversion consists of a series of biologically plausible, homogeneous transformations based on Predictive Coding principles. Under the plausible assumption of a nonuniform fovea resolution, depth estimation favors an active vision strategy that fixates the object with the eyes, rendering the depth belief more accurate. This strategy is not realized by first fixating on a target and then estimating the depth, but by combining the two processes through action-perception cycles, with a similar mechanism of the saccades during object recognition. The proposed approach requires only local (top-down and bottom-up) message passing that can be implemented in biologically plausible neural circuits.

## 1 Introduction

Depth estimation is a complex process involving continuous activation of every level of the visual cortex and even higher-level regions. Disparity-sensitive cells of a different kind can be found early in the visual cortex [1, 2], and it seems that the resulting signals travel through the dorsal and ventral pathways for different purposes: parietal regions (in particular anterior and lateral intraparietal) have major contributions in depth estimation for visually-guided actions in hand and eye movements [3, 4], while the inferotemporal cortex supports the creation of 3D shapes based on the relative disparity between objects [2, 5]. The brain can rely on several cues to estimate the depth of objects, the most important ones being: (i) the binocular disparity that allows the visual cortex to have access to two different perspectives of the same environment; (ii) the motion parallax effect that happens when objects at a greater distance move slower than nearby objects; (iii) the angular difference between the eyes when fixating the same object (*vergence*). However, the exact contribution of these mechanisms on the overall depth estimation and, critically, where and how the information processing of these signals occurs, is still unclear.

Traditionally, the visual cortex has been associated with a feature detector: as the sensory signals climb the hierarchy, more complex features are detected by increasing cortical levels, such that from lines and contours high-level representations of objects are constructed. This view has inspired the development of Convolutional Neural Networks, which have led to remarkable results in object recognition tasks [6]. Despite its success, this bottom-up approach is not able to capture several top-down mechanisms that affect our everyday perception of the external world, as in the case of visual illusions [7]. In recent years, a different perspective has emerged, based on Predictive Coding theories, that views these illusions not just as unexpected phenomena but as expressions of the main mechanism through which our brain is able to efficiently predict and act over the environment [8, 9]. Under this view, the biases we perceive are actually hints to better minimize the errors between our sensations and our predictions [10]. Furthermore, vision is increasingly considered to be an active process that constantly tries to reduce the uncertainty of what will happen next.

In this article, we apply this predictive and inferential view of perception to depth estimation. Specifically, we advance an Active Inference model that is able to estimate the depth of an object based on two projected images, through a process of prediction error minimization and active oculomotor behavior. In our model, depth estimation does not consist of a bottom-up process that detects disparities in the images of the two eyes, but of an inference of top-down projective predictions from a high-level representation of the object. In other words, the estimation of the object’s depth naturally arises by inverting a visual generative model where the resulting prediction errors flow up the cortical hierarchy, in contrast with the direct process occurring with neural networks.

## 2 Methods

The theory of Active Inference assumes that an agent is endowed with a generative model that makes predictions over sensory observations [10, 11, 12, 13], as shown in Figure 1. The discrepancy between predictions and observations generates a prediction error that is minimized in order to deal with a dynamical environment and to anticipate what will happen next. This generative model hinges on three components encoded in generalized coordinates of increasing temporal orders (e.g., position, velocity, acceleration, and so on): hidden states 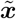, hidden causes 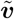, and sensory signals 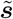. These components are expressed through a non-linear system that defines the prediction of sensory signals and the evolution of hidden states and causes across time:

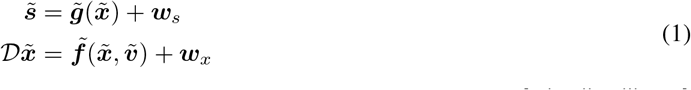

**Figure 1.**
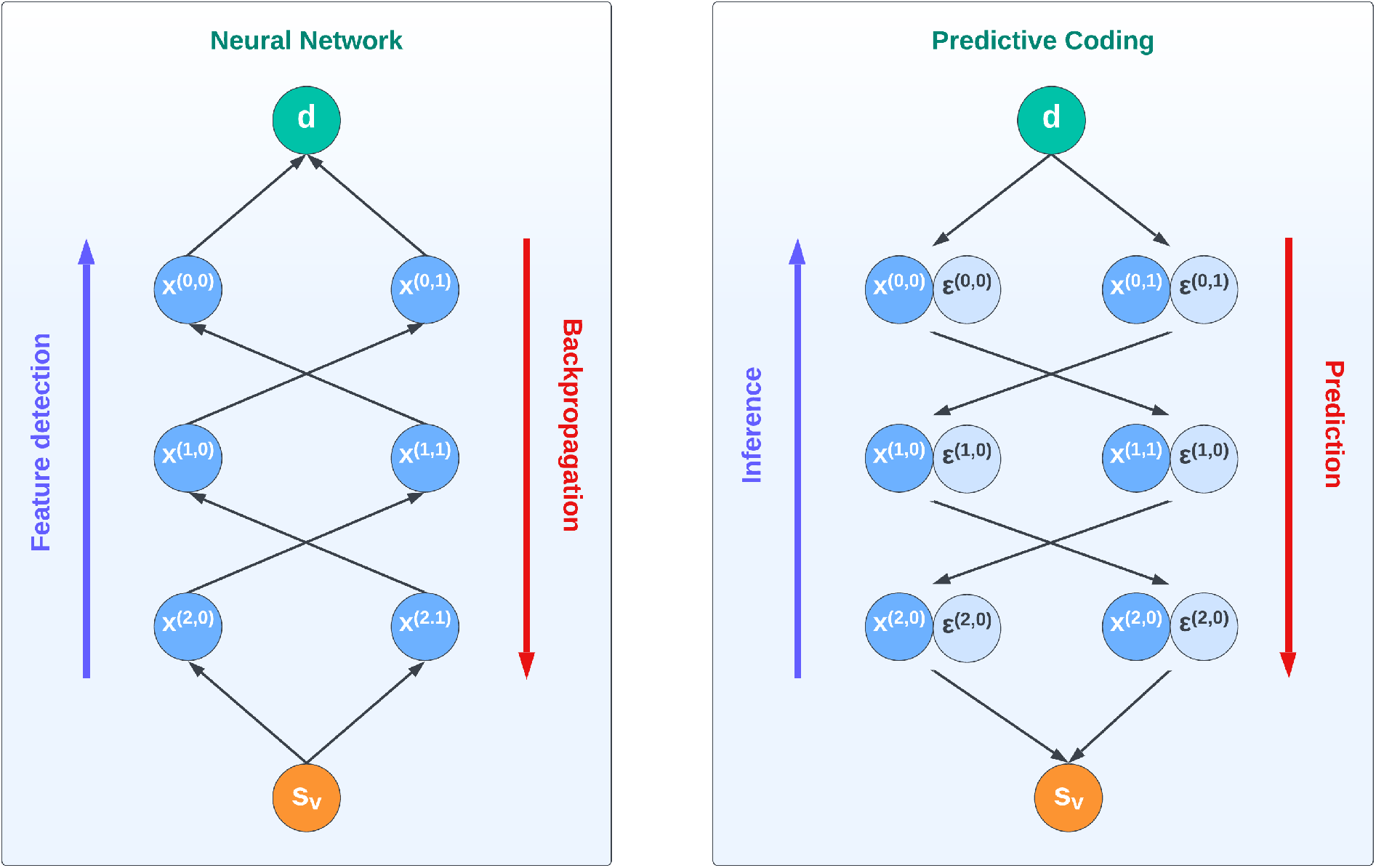
Information processing in neural networks (left) and Predictive Coding (right). In a neural network, the visual observation **s**_*v*_ travels through the cortical hierarchy in a bottom-up way, detecting increasingly more complex features ***x***^(*i,j*)^ and eventually estimating the depth of an object ***d***. The descending projections are considered here as feedback signals that convey backpropagation errors. In contrast, in a Predictive Coding Network the depth ***d*** is a high-level belief generating a visual prediction that is compared with the observation. This process leads to a cascade of prediction errors ***ε***^(*i,j*)^ – associated with each intermediate prediction ***x***^(*i,j*)^ – that are minimized throughout the hierarchy, eventually inferring the correct belief (for details, see the Methods section).

In this context, 𝒟denotes a differential operator that shifts all temporal orders by one, such as 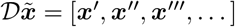. Furthermore, ***w***_*s*_ and ***w***_*x*_ stand as noise terms drawn from a Gaussian distribution. The considered joint probability is divided into distinct distributions:

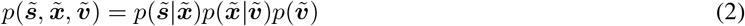

Typically, each distribution is approximated with Gaussian functions:

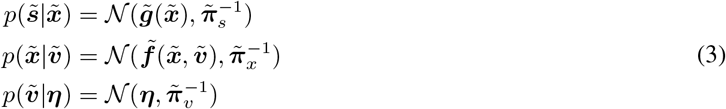

where ***η*** is a prior, and the distributions are expressed in terms of precisions (or inverse variances) 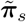,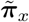, and 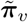.

Following a variational inference method [14], these distributions are inferred via approximate posteriors 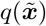 and 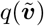. Under appropriate assumptions, minimizing the Variational Free Energy (VFE) ℱ, defined as the disparity between the KL divergence of real and approximate posteriors and the log evidence:

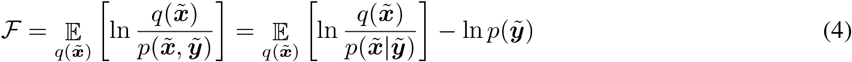

leads to the minimization of prediction errors. The belief updates 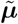 and 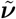, concerning hidden states and hidden causes respectively, expand as follows:

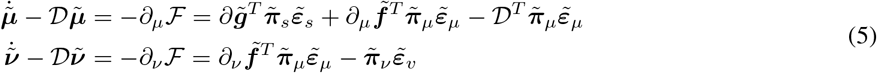

Here,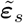,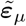, and 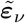 denote the prediction errors of sensory signals, dynamics, and priors:

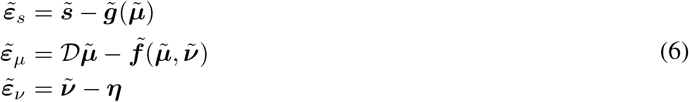

A simple Active Inference scheme can handle various tasks, yet the effectiveness of the theory stems from a hierarchical structure that enables the brain to grasp the hierarchical associations between sensory observations and their causes [15]. Specifically, the model delineated above can be expanded by linking each hidden cause with another generative model; as a result, the prior becomes the prediction from the layer above, while the observation becomes the likelihood of the layer below:

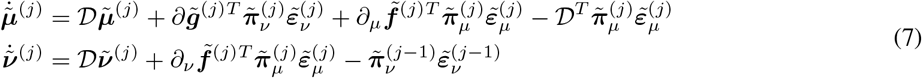

Here:

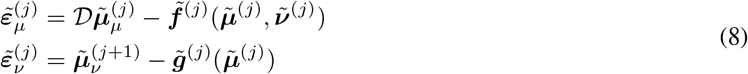

In contrast, the execution of action is accomplished by minimizing the proprioceptive component of the VFE concerning the motor control signals ***a***:

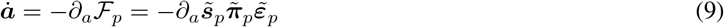

Here, *∂*_*a*_***s***_*p*_ stands for the partial derivative of proprioceptive observations regarding the motor control signals,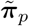 are the precisions of the proprioceptive generative model, and 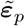 are the generalized proprioceptive prediction errors:

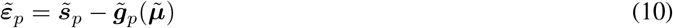

In summary, in Active Inference goal-directed behavior is generally possible by firstly biasing the belief over the hidden states through a specific cause. The latter acts as a prior that encodes the agent’s belief about the state of affairs of the world. In this context, action follows because the hidden states generate a proprioceptive prediction error that is suppressed through a reflex arc [16]. For instance, suppose that the agent has to rotate the arm by a few degrees. The belief over its arm’s angle is subject to two opposing forces:

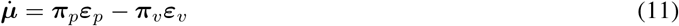

one coming from above (pulling it toward its expectation) and one coming from below (pulling it toward what it is currently perceiving). The tradeoff between the two forces is expressed in terms of the precisions ***π***_*p*_ and ***π***_*v*_, which encode the agent’s level of confidence about the particular prediction errors. By appropriately tuning the precision parameters, it is possible to smoothly push the belief toward a desired state, eventually driving the real arm through Equation 9.

## 3 Results

### 3.1 Homogeneous transformations as Hierarchical Active Inference

Classical Predictive Coding models are passive, in the sense that the model cannot select its visual stimuli [8]. Rather, our Active Inference model can actively control “eyes”, to sample the preferred stimuli that reduce prediction errors.

State-of-the-art implementations of oculomotor behavior in Active Inference rely on a latent state (or belief) over the eye angle, and attractors are usually defined directly in the polar domain [17, 18]. While having interesting implications for simulating saccadic and smooth pursuit eye movements, such models do not consider the fact that eyes fixate the target from two different perspectives. A similar limitation can be found in models of reaching, in which the 3D position of the object to reach is directly provided as a visual observation [19, 20]. Furthermore, since there is only a single level specified in polar coordinates, if one wants to fixate or reach a target defined in Cartesian coordinates, a relatively complex dynamics function has to be defined at that level.

Using a *hierarchical kinematic model* – based on Active Inference – that includes both intrinsic (e.g., joint angles and limb lengths) and extrinsic (e.g., Cartesian positions) coordinates affords efficient control of a simulated body [21]. The extrinsic configuration of the motor plant is computed hierarchically, as shown in Figure 2. For the relatively simple kinematic control tasks targeted in [21], these computations only required two simple transformations between reference frames: translations and rotations. However, a hierarchical kinematic model can be easily extended to afford more complex tasks, requiring different transformations.

**Figure 2.**
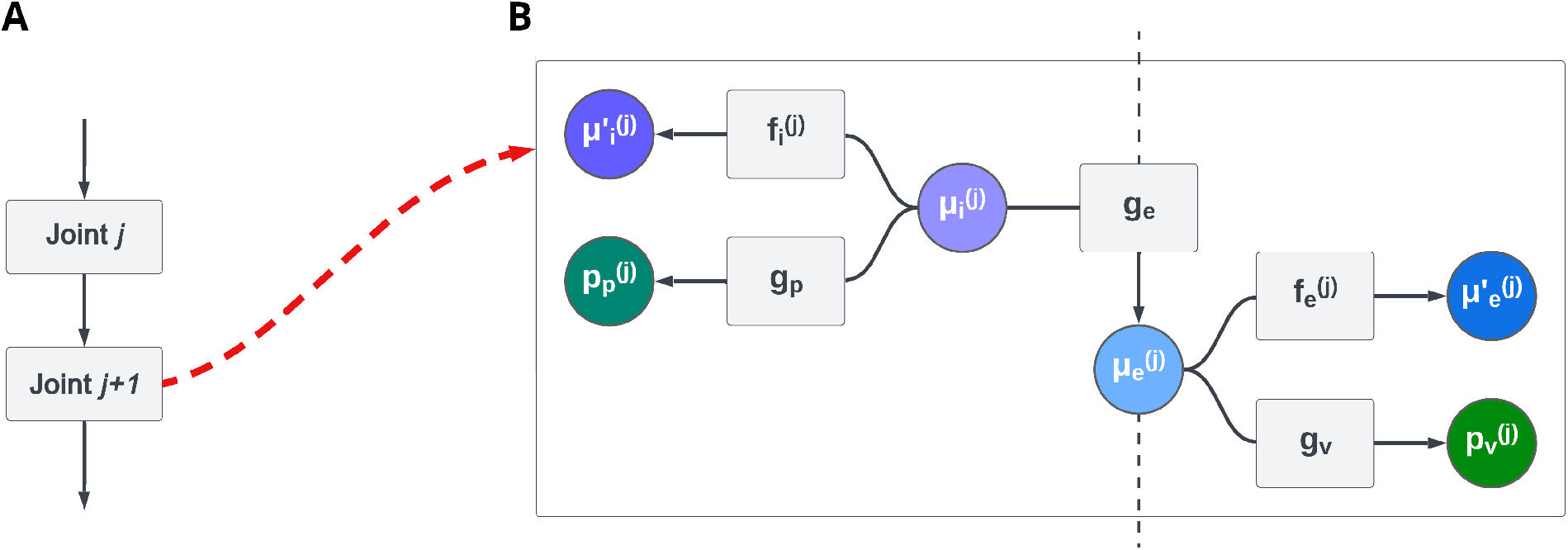
(**A**) An example of a portion of a kinematic plant. (**B**) Factor graph of a single level *j* of the hierarchical kinematic model composed of intrinsic 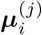 and extrinsic 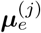 beliefs. These beliefs generate proprioceptive and visual predictions 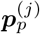 and 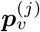 through generative models ***g***_*p*_ and ***g***_*v*_, respectively. Furthermore, the beliefs predict trajectories – here, only velocities 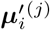 and 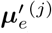 – through dynamics functions 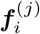 and 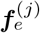. Note that the extrinsic belief of level *j −*1 acts as a prior for layer *j* through a kinematic generative model ***g***_*e*_. See [21] for more details.

In robotics, transformations between reference frames are usually realized through the multiplication of a linear transformation matrix. These operations can be decomposed into simpler steps where *homogeneous coordinates* are multiplied once at a time through the chain rule, allowing more efficient computations. Specifically, if the x and y axes represent a Cartesian plane, a homogeneous representation augments the latter with an additional dimension called *projective space*. In this new system, multiplying the point coordinates by the same factor keeps the mapping unaltered, i.e., (*p*_*x*_, *p*_*y*_, 1) *≡* (*p*_*z*_*p*_*x*_, *p*_*z*_*p*_*y*_, *p*_*z*_).

Affine transformations preserve parallel lines and have the following form:

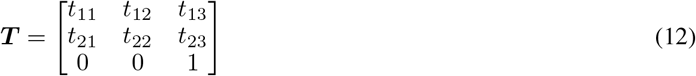

where the last row ensures that every point always maps to the same plane. By the chain rule, a point in the plane ***p***^0^ can be rotated and translated by multiplication of the corresponding transformations:

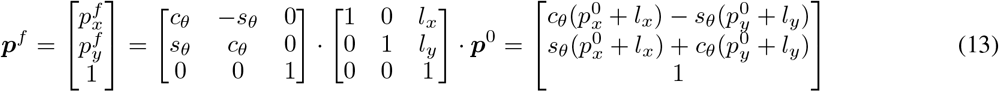

where *s*_*θ*_ and *c*_*θ*_ are the sine and cosine of the rotation *θ*, while *l*_*x*_ and *l*_*y*_ are the coordinates of the translation.

By appropriately changing the values of the matrix, more affine transformations can be obtained, such as shearing or scaling. Critically, if the last row is modified, it is also possible to realize perspective projections:

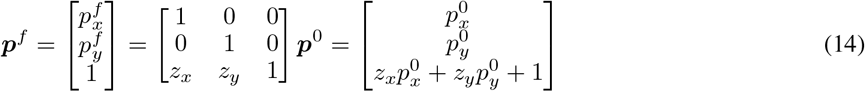

so that the new point does not map on the same plane *p*_*z*_ = 1 as before. Thus, to map it back to the Cartesian plane, we divide the x and y coordinates by the last element:

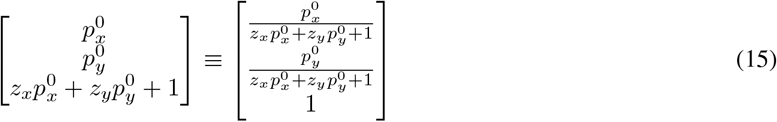

This special transformation is critical for computer vision in order to project points on an image plane or to estimate the depth of an object. If we have a 3D point ***a*** = (*a*_*x*_, *a*_*y*_, *a*_*z*_, 1) expressed in homogeneous coordinates, we can obtain the corresponding 2D point ***p*** projected in the camera plane by first performing a roto-translation – similar to Equation 13 – through a matrix that encodes where the camera is located and oriented (the *extrinsic parameters*):

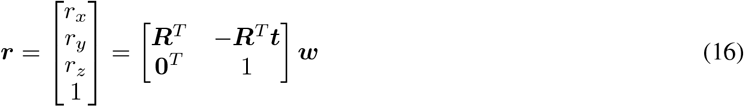

and then scaling and converting the point to 2D through the so-called *camera matrix*:

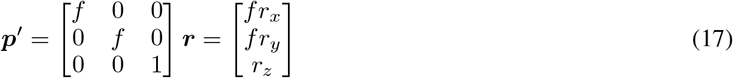

The projection is performed by multiplying the depth coordinate z by the focal length *f*, which represents the distance of the image plane from the origin. As before, since the homogeneous representation is up to a scale factor, to transform the point ***p***^*′*^ into the Cartesian space we divide the camera coordinates by the depth coordinate *p′*_*z*_ (as shown in Figure 3):

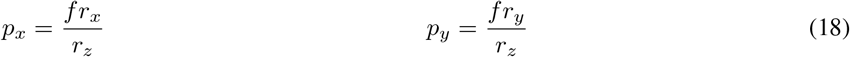

**Figure 3.**
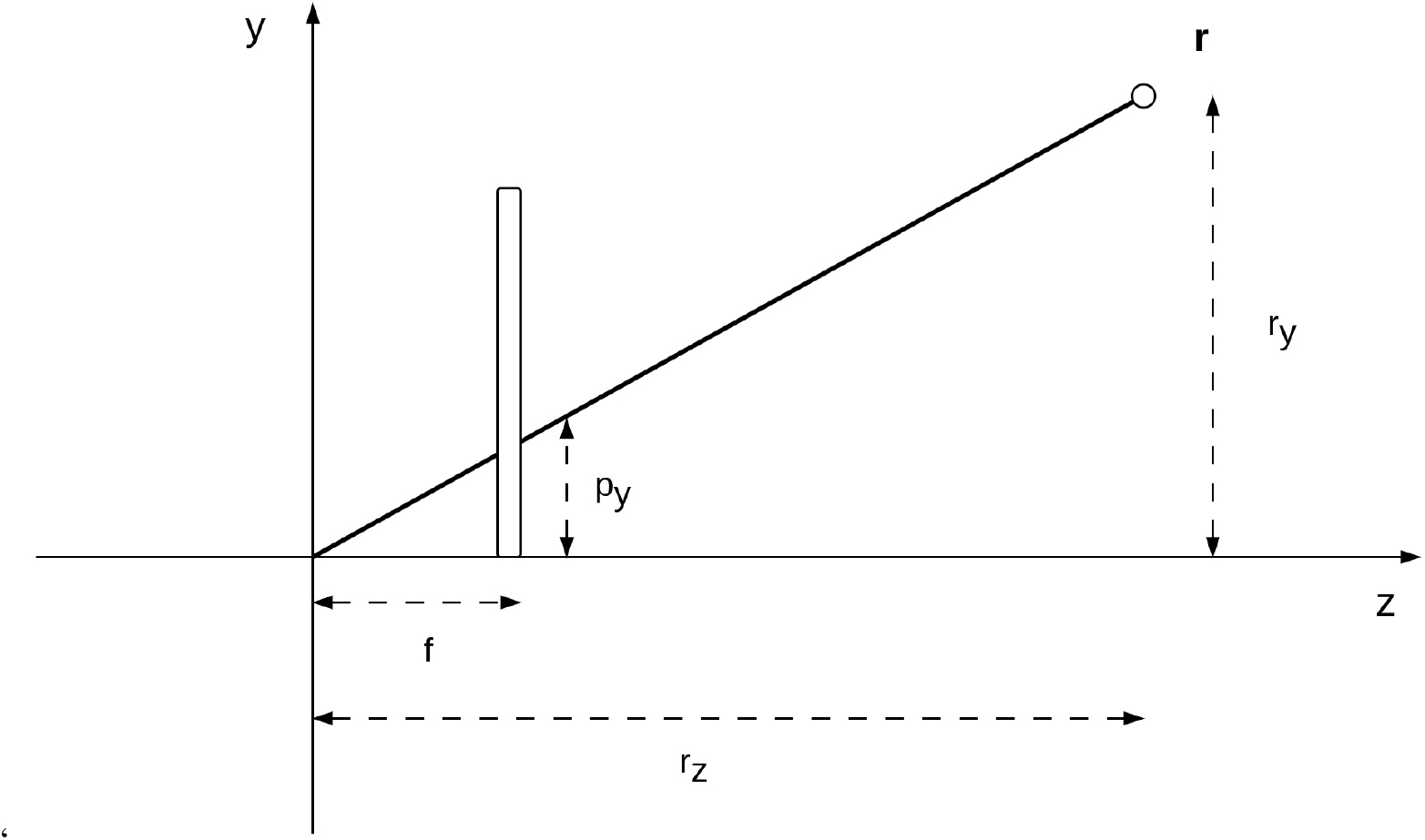
Projection of a 3D point in the camera plane (only two dimensions are shown). The y coordinate of the real point *r*_*y*_ and of the projected point *p*_*y*_ are related by the ratio between the focal length *f* and the real point depth *r*_*z*_.

Keeping this in mind, we generalize the deep kinematic model of [21] by assuming that each level sequentially applies a series of homogeneous transformations. Specifically, the first belief, called 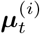, contains information about a particular transformation (e.g., by which angle to rotate or by which length to translate a point).

This belief then generates a homogeneous transformation relative to that Degree of Freedom (DoF), which is then multiplied by a second belief expressed in a particular reference frame (as exemplified in Figure 4):

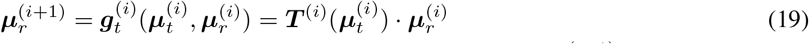

**Figure 4.**
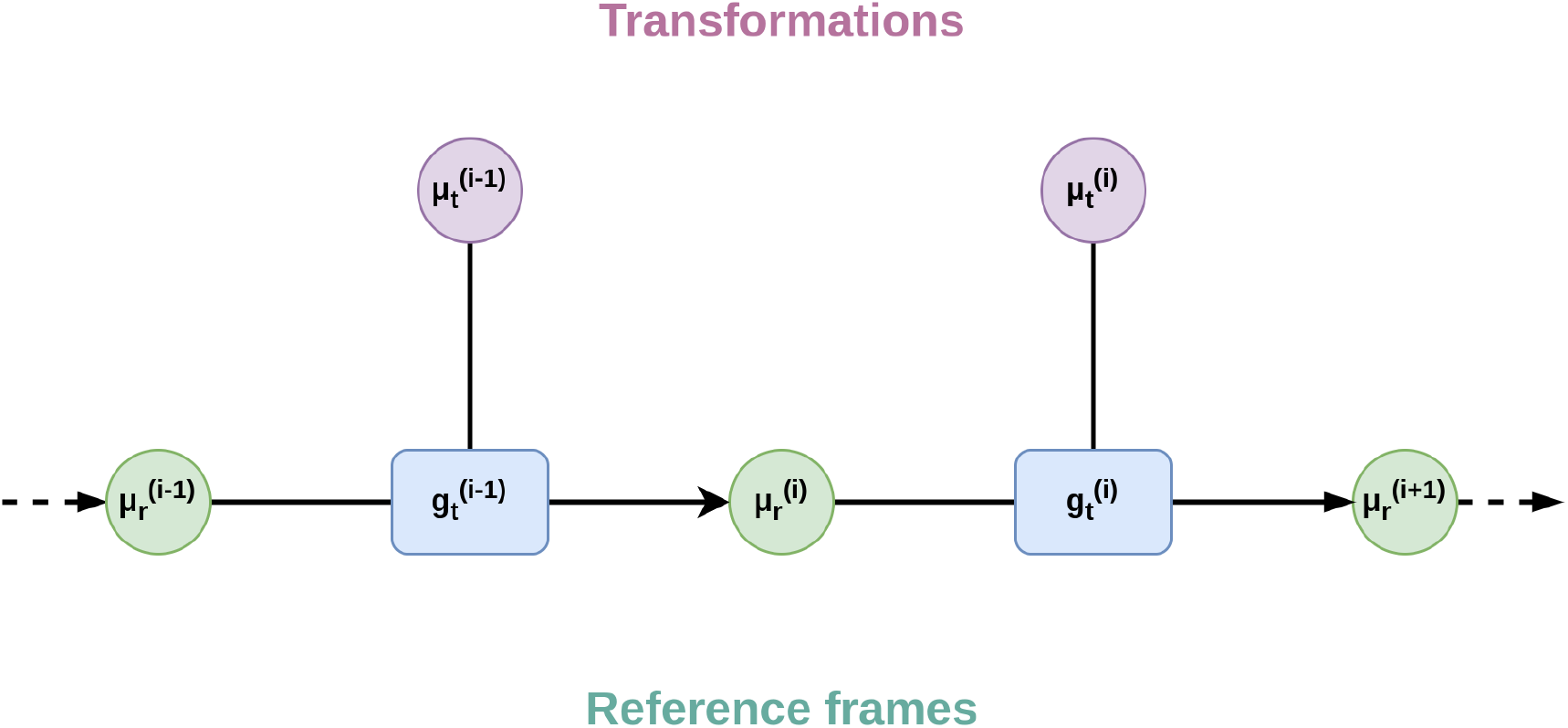
Representation of the hierarchical relationships of a generalized model with homogeneous transformations. The belief over a reference frame ***μ***_*r*_ of level *i* is passed to a function ***g***_*t*_ encoding a homogeneous transformation along with a belief over a particular transform ***μ***_*t*_ (e.g., angle for rotation or length for translation), generating the reference frame of level *i* + 1.

This equation leads to simple gradient computations through the generated prediction error 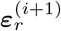 iteratively update the two beliefs:

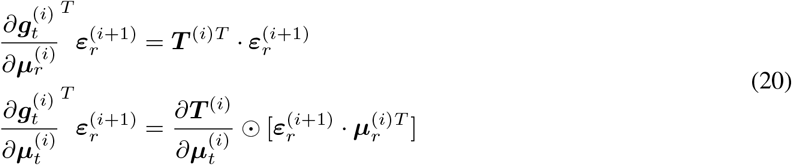

where ⊙ is the element-wise product.

### 3.2 A hierarchical generative model for binocular depth estimation

In this section, we explain how depth estimation arises by inverting the projective predictions of the two eyes, using a hierarchical generative model. For simplicity, we consider an agent interacting with a 2D world, where the depth is the *x* coordinate. Nonetheless, the same approach could be used to estimate the depth of a 3D object. We construct the generative model hierarchically, starting from a belief ***μ***_*a*_ = (*μ*_*a,x*_, *μ*_*a,y*_, 1) about the absolute 2D position of an object encoded in homogeneous coordinates, where *μ*_*a,x*_ is the depth belief. Then, two parallel pathways generate specular predictions 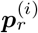 that receive the eye angles encoded in a common vergence-accommodation belief ***μ*** = (*θ, θ*) – and transform the absolute coordinates of the object into the two reference frames relative to the eyes:

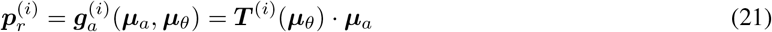

where ***T***^(*i*)^(***μ***_*θ*_) is the homogeneous transformation corresponding to the extrinsic parameters of the camera:

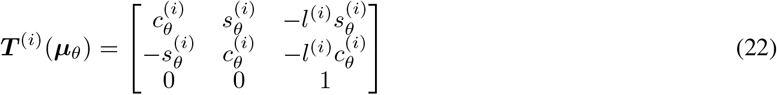

Here, *l*^(*i*)^ is the distance between an eye and the origin (i.e., the middle of the eyes), and the absolute eye angles are:

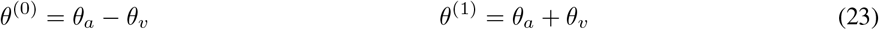

Each of these beliefs generates a prediction over a point projected to the corresponding camera plane:

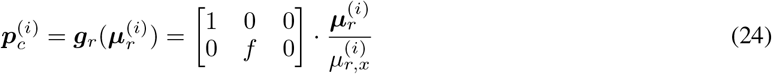

Figure 5 provides a neural-level illustration of the model, with the two branches originating from the two beliefs at the top. Note that while the eye angles belief ***μ***_*θ*_ generates separate predictions for the two eyes, proprioceptive predictions directly encode angles in the vergence-accommodation system, which is used for action execution [10].

**Figure 5.**
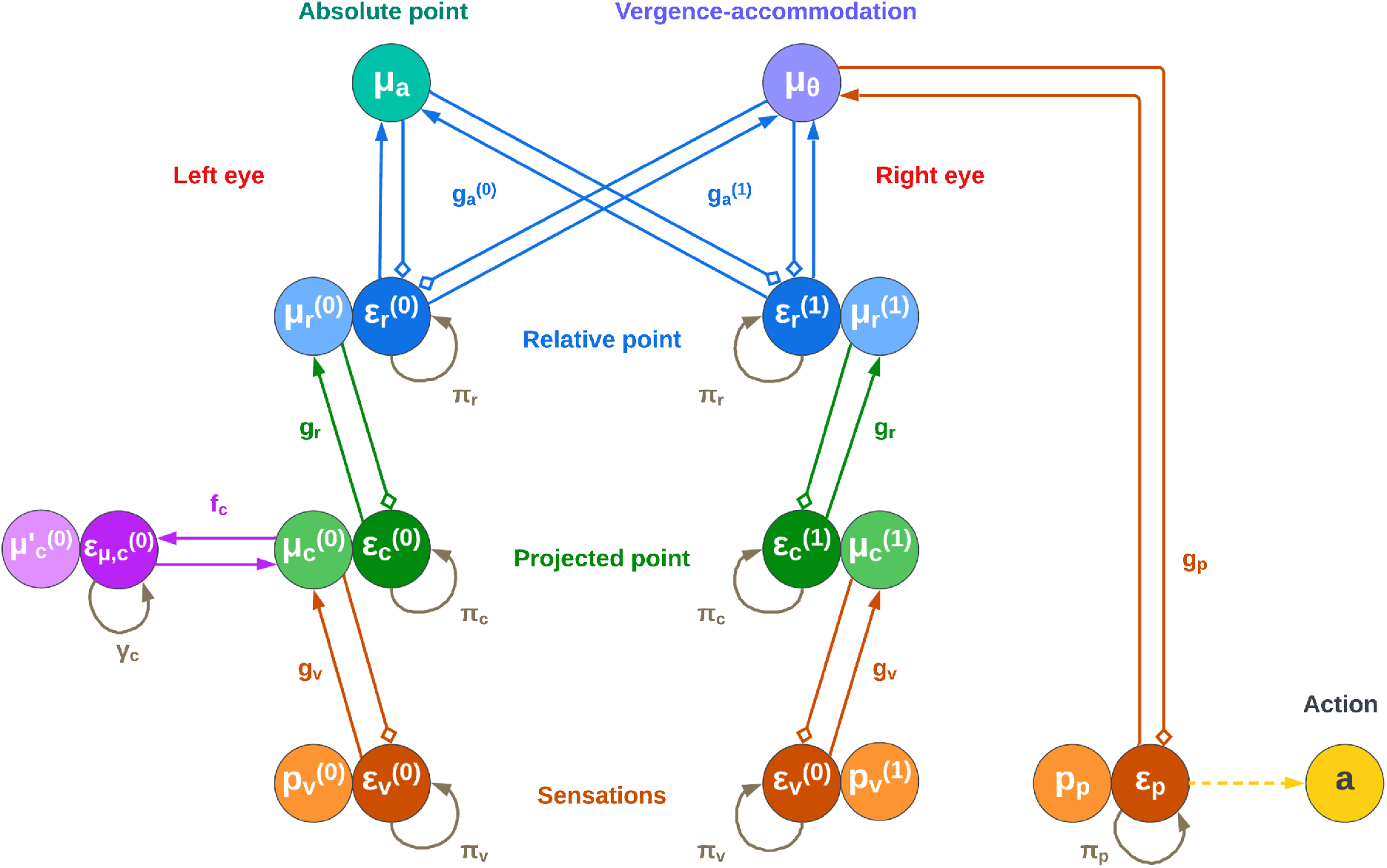
Neural-level implementation of a hierarchical generative model to estimate the depth of a point, through Active Inference. Small squares indicate inhibitory connections. Differently from a neural network, depth is estimated by first generating, from a point in absolute coordinates ***μ***_*a*_ and the vergence-accommodation angles ***μ***_*θ*_, two predictions 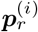 of the point relative to each eye. The new belief in turn computes a projection 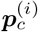, and finally a visual prediction 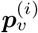. The predictions are then compared with the visual observations, generating prediction errors throughout the hierarchy and eventually driving the beliefs at the top toward the correct values. Note that eye movements are directly triggered to suppress the proprioceptive prediction error ***ε***_*p*_. Intentional eye movements (e.g., for target fixation) can be instead achieved by setting a prior in the dynamics function ***f***_*c*_ of the belief over the projected point ***μ***_*c*_ (note that for better readability, the figure only shows the dynamics function ***f***_*c*_.

The absolute point belief (encoded in generalized coordinates up to the 2nd level) is updated as follows:

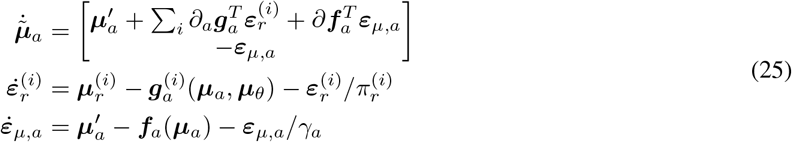

where 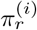 and 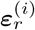 are the precisions and prediction errors of the beliefs below, while ***f***_*a*_, *γ*_*a*_ and ***ε***_*μ,a*_ are the function, precision, and prediction error of the dynamics of the same belief.

This belief is thus subject to different prediction errors 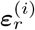 coming from the two eyes, and the depth of the point is estimated by averaging these two pathways. Furthermore, an attractor can be defined in the dynamics function ***f***_*a*_ if one wants to control the object encoded in absolute coordinates – e.g., for reaching or grasping tasks.

Similarly, the belief update equation for ***μ***_*θ*_ is the following:

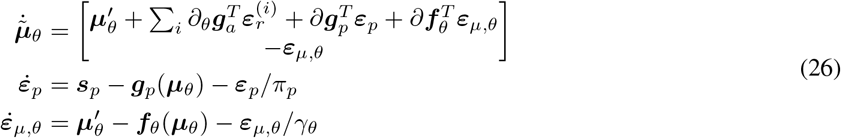

where *π*_*p*_, ***ε***_*p*_, ***s***_*p*_, and ***g***_*p*_ are the proprioceptive precision, prediction error, observation, and likelihood function – which is an identity mapping in the following simulations – while ***f***_*θ*_, *γ*_*θ*_ and ***ε***_*μ,θ*_ are the function, precision, and prediction error of the belief dynamics.

This belief, besides being affected by the proprioceptive contribution, is also subject to the same prediction errors 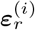 in the same way as the absolute belief. The overall free energy can be thus minimized along two different pathways: (i) by changing the belief about the absolute location of the object (including the depth), or (ii) by modifying the angle of fixation of the eyes. As will be shown in the next section, the possibility of using both pathways may create some stability issues during goal-directed movements.

Also in this case, an attractor can be specified in the dynamics function ***f***_*θ*_ if one wants to explicitly control the dynamics of the eyes – e.g., not fixating a point on the camera plane, but rotating the eyes along a particular direction or by a particular angle.

Finally, the belief update equation for the projected point 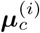 is the following:

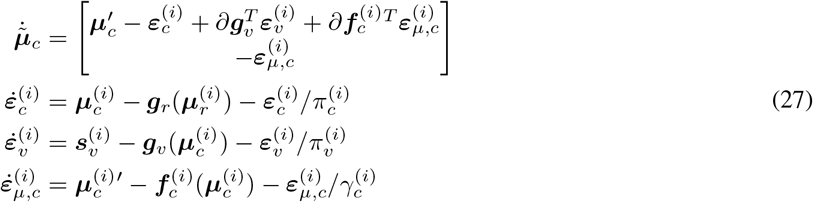

where 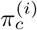 and 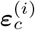 are the precisions and prediction errors of the beliefs below,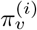,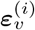,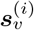,and ***g***_*v*_are the visual precision, prediction error, observation, and likelihood function, while ***f***_*c*_, *γ*_*c*_ and ***ε***_*μ,c*_ are the function, precision, and prediction error of the belief dynamics. Note that in the following simulations, we approximated ***g***_*v*_ by a simple identity mapping, so that 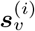 conveys a Cartesian position.

Differently from the belief over the eye angles ***μ***_*θ*_ which is only biased by the likelihoods of the levels below, this belief is both subject to a prior encoded in 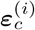, and a visual likelihood from 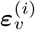.

### 3.3 Active vision and target fixation with action-perception cycles

The model advanced here can not only infer the depth of a point, but also fixate it using active vision. This is possible by specifying an appropriate attractor in the dynamics function of the last belief 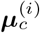, or in other words, by setting an “intention” in both eyes [22], such that the projected position will be at the center of the camera planes:

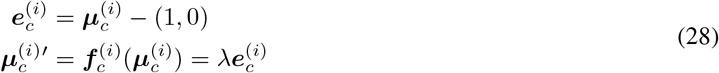

In short,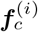 returns a velocity encoding the difference 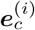 between the current belief and the center of the camera plane – expressed in homogeneous coordinates – multiplied by an attractor gain *λ*. Thus, the agent thinks that the projected point will be pulled toward the center, with a velocity proportional to 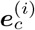. The prediction errors generated then travel back through the hierarchy, affecting both the absolute and the eye angles beliefs. Since what we want in this case is modifying the latter (directly generating proprioceptive predictions for movement), the former pathway can be problematic. In fact, if ***μ***_*a*_ already encodes the correct depth of the object, fixation occurs very rapidly; however, if this is not the case, the prediction errors 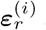 are free to flow through all the open pathways, driving the beliefs toward different directions and leading the free energy minimization process to be stuck with incorrect depth coordinate and eye angles [23].

This abnormal behavior can be avoided if we decompose the task into cyclical phases of action and perception [24]. During an action phase, the absolute belief is kept fixed, so that the relative prediction errors 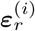 can only flow toward the belief over the eye angles, resulting in the eyes moving according to the depth belief. During a perception phase, action is blocked (e.g., either by setting the attractor gain or the proprioceptive precision to zero), while 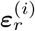 is free to flow in any direction, which has the result of pushing the depth belief toward the correct value signaled through the sensory observations. Depth estimation is therefore achieved through multiple action and perception cycles until the overall free energy is minimized.

Figure 6 shows a sequence of time frames of a depth estimation task, in which the (perceptual) process of depth estimation and the (active) process of target fixation cyclically alternate in different phases every 100 time steps. As can be seen from the visualization of the point projections, the distance between the real and estimated target positions slowly decreases, while both positions approach the center of the camera planes, affording efficient depth estimation.

**Figure 6.**
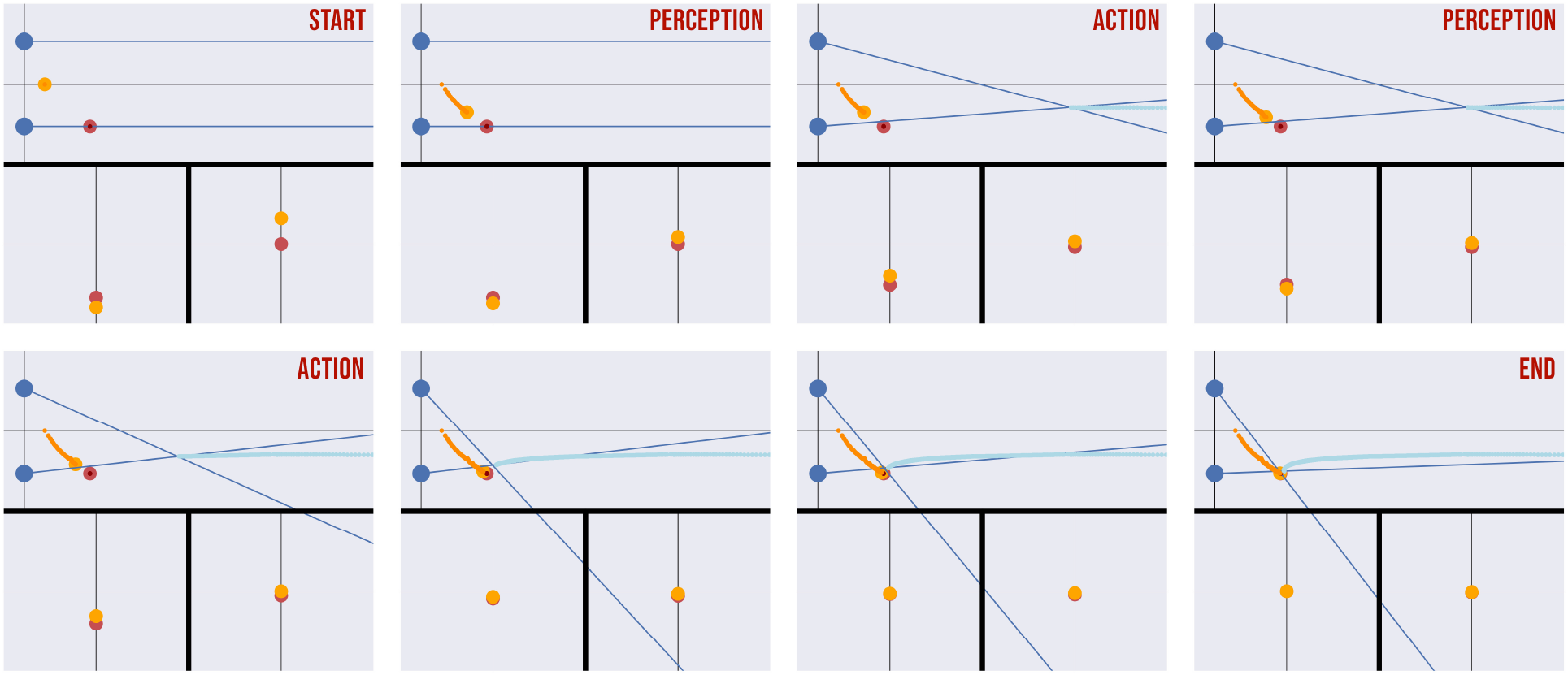
Sequence of time frames of a depth estimation task with simultaneous target fixation. The agent uses alternating action-perception phases to avoid being stuck during the minimization process. Each frame is composed of three images: a third-view perspective of the overall task (top), and a first-view perspective consisting of the projection of the target to the camera planes of both eyes (bottom left and bottom right). In the top panel, the eyes are represented with blue circles, while the real and estimated target positions are in red and orange. The fixation trajectory (when vergence occurs) is represented in cyan. The thin blue lines are the fixation angles of the eyes. In the bottom panel, the real and estimated target positions are in red and orange. The abscissa and the ordinate represent target depth and its projection.

### 3.4 Model comparison

We tested the model introduced in sections 3.2 and 3.3 in a depth estimation task that consists of inferring the 2D position of the object shown in Figure 6. We compared three different versions of the model. In the first version, the eyes are kept in a fixed position, with the eyes parallel to one another (*infer parallel*). In the second and third versions, the model can actively control the eye angles, but the initial values are set at the correct target position (*infer vergence*) or at a random location (*active vision*, as displayed in Figure 6).

Furthermore, the fovea of the simulated “eye” could have either a uniform or a nonuniform resolution, where in the latter condition the object is represented with greater accuracy when it is near the point of fixation. This reflects the fact that the biological fovea has far more receptors at the center than in the peripheral vision, which has been modeled with an exponential link [25]. Specifically, in our implementation the variability **Σ**_*v*_ of the Gaussian error in the visual observations (i.e., in the generative process) exponentially increases with the distance *d* between the point of fixation and the real target position:

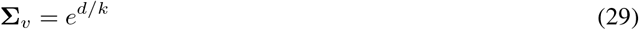

where *k* is a scaling factor (equal to 1.5 in the simulations). In the uniform condition, the visual noise was set to zero.

Figure 7 shows the results of the simulations, including accuracy (the number of trials in which the agent successfully predicts the 2D position of the target, left panel), mean error (distance between the real and estimated target position at the end of every trial, middle panel), and estimation time (number of steps needed to correctly estimate the target, right panel). The number of time steps for each phase has been set to 100, as before. The figure shows that depth estimation with parallel eyes (*infer parallel*) in the nonuniform condition results in a very low accuracy, especially when the target position is far from the fixation point. This is to be expected, since in this condition the fovea has low resolution at the periphery. Rather, if the angle of the eyes is set to fixate on the target (*infer vergence*), the accuracy is much higher and few fluctuations occur. Finally, the *active vision* model that simultaneously implements depth estimation and target fixation achieves a level of performance that is almost on par with the model where the fixation is initialized at the correct position. Indeed, the only appreciable difference between the last two conditions is the slightly greater number of time steps in the *active vision* condition.

**Figure 7.**
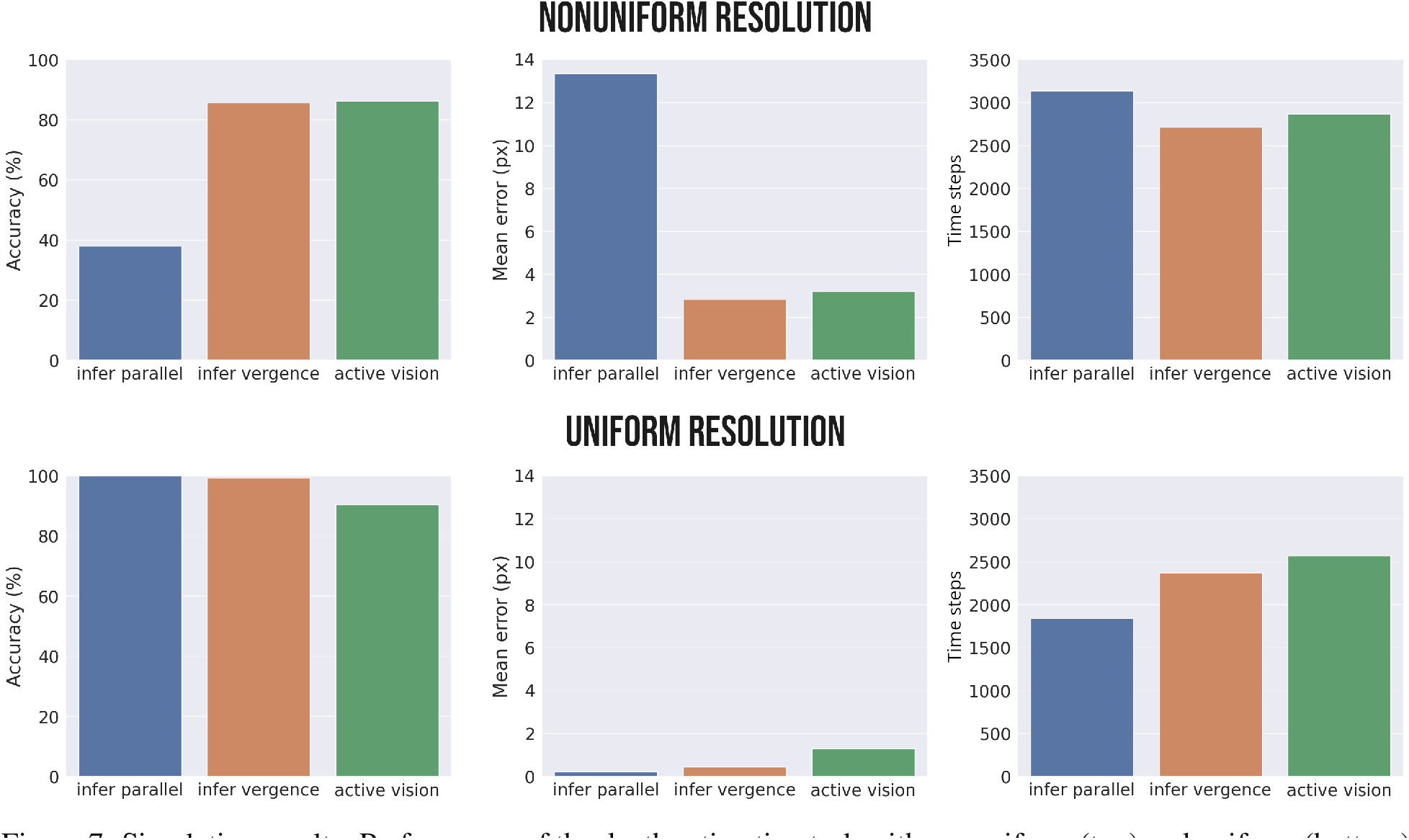
Simulation results. Performance of the depth estimation task with nonuniform (top) and uniform (bottom) fovea resolution during inference with the eyes parallel and fixed (*infer parallel*), inference with the eyes fixating on the target (*infer vergence*), and simultaneous inference and target fixation (*active vision*). The accuracy (left panel) measures the number of trials in which the agent successfully predicts the 2D position of the target, the mean error (middle panel) measures the distance between the real and estimated target positions at the end of every trial, and the time (right panel) measures the number of steps needed to correctly estimate the target.

This pattern of results shows three main things. First, the hierarchical Active Inference model is able to solve the depth estimation problem, as evident from its perfect accuracy in the task. Second, the model is able not only to infer depth but also to simultaneously select the best way to sample its preferred stimuli (i.e., by fixating on the target). This is possible because, during a trial (as exemplified graphically in Figure 6), the active vision model obtains increasingly more accurate estimates of the depth, as the point of fixation approaches the target. Note that this pattern of results emerges because of the nonuniform resolution of the fovea. In fact, if the fovea resolution is assumed to be uniform (such as in camera models of artificial agents), the best accuracy is achieved by keeping the eyes parallel (Figure 7, *infer parallel* condition). In this case, fixating the target does not help but rather hinders and slows down depth estimation, which is probably caused by the increased effort that the agent needs to make in order to infer the reference frames of the eyes rotated in different directions. This has the consequence of further increasing the time needed to estimate the depth for the active vision model.

Intuitively, the better performance in the uniform condition is due to the lack of noise in the visual input. Although a more realistic scenario should consider noise also in this case, it is reasonable to assume that it would have a much smaller amplitude, due to a uniform distribution. Considering the nonuniform sensory distribution only, the better performance in the *infer vergence* condition relative to *active vision* could be due to the fact that in the former case the agent starts the inference process from a fixation on the correct 3D position. Thus, an active vision strategy in the *infer vergence* condition only needs to estimate the object’s depth. If we compare these two scenarios, we can note that *active vision* performs almost optimally, similar to the *infer vergence* condition when the eye angles are set to the correct values for object fixation. However, the latter condition rarely occurs in a realistic setting and a more meaningful depth estimation comparison would be between *active vision* and the more general case in which the agent is fixating another object, or not fixating anything in particular – which we approximated with the *infer parallel* simulation.

## 4 Discussion

We advanced a hierarchical Active Inference model for depth estimation and target fixation operating in the state-space of the projected camera planes. Our results show that depth estimation can be solved by inference – that is, by inverting a hierarchical generative model that predicts the projection of the eyes from a 2D belief over an object. Furthermore, our results show that active vision dynamics makes the inference particularly effective – and that fixating the target drastically improves task accuracy (see Figure 7). Crucially, the proposed model can be implemented in biologically plausible neural circuits for Predictive Coding [8, 9, 10], which only require local (top-down and bottom-up) message passing. From a technical perspective, our model shows that inference in any homogeneous transformation can be iteratively realized by combining generative models at different levels, each of which computes a specific transformation that can be, e.g., a roto-translation for kinematic inference [21], or a projection for computer vision.

This proposal has several elements of novelties compared to previous approaches to depth estimation. First, by focusing on inference and local message passing, our proposal departs from the trend of viewing cortical processing from a purely bottom-up perspective. The latter is common in neural network approaches, which start from the image of an object and gradually detect more and more complex features, eventually estimating its depth. Our proposal is also distinct from a direct approach that generates the depth of an object from a top-down perspective, e.g., with vergence cues. The role of vergence has been considered key in facilitating binocular fusion [5] and maximizing coding efficiency in a single environmental representation [26], but recent studies have dramatically reduced the importance of this mechanism in depth estimation. Binocular fusion might not be strictly necessary for this task [27] – 90% performance of depth estimation is attributed to diplopia [28] – and has never actually been tested as an absolute distance cue without eliminating all possible confounds [28]. Moreover, when fixating a target, there is always a vertical fixation disparity of monocular images, with no line precisely intersecting to form a vergence angle [29] – and it has been demonstrated that vergence does not correspond to the exact distance of the object being gazed at [30]. In keeping with this body of evidence, in our model the vergence belief does not play a critical role in depth estimation: it only operates along with a high-level belief over the 2D position of the object in order to predict the projections for the two eyes. These projections are compared with the visual observations, and the resulting prediction errors that flow back through the hierarchy drive the update of both beliefs (about the eye angles and the 2D object position), in two possible ways: they might change (i) the estimated depth of the object, or (ii) the vergence-accommodation angles of the eyes, ultimately realizing a specific movement. In sum, depth estimation is not purely a top-down process, but rather it is realized through the inversion of a generative projective model and by averaging the information obtained through the two parallel pathways of the eyes. In conclusion, our model supports the direct (from disparity to both vergence and depth) rather than the indirect (from disparity to vergence, and then to depth) hypothesis of depth estimation [27]. Furthermore, this account is in line with the fact that small changes in vergence (*delta theta*) are the consequence and not a direct hint about depth estimation [28], and that reflex-like vergence mechanisms serve only to eliminate small vergence errors and not to actively transfer the gaze to new depth planes [31].

Second, an interesting consequence of this architecture is that – in contrast to standard neural networks – it permits imposing priors over the depth belief in order to drive and speed up the inferential process. Such priors may come through different sensory modalities or other visual cues, e.g., motion parallax or perspective, which we did not consider here. This is in line with the finding that vergence alone is unable to predict depth with ambiguous cues [5], suggesting that depth belief is constantly influenced by top-down mechanisms and higher-level cues, and not just directly from perception. Besides depth priors, using an Active Inference model has also the advantage that, if one wants to fixate on a target, simple attractors can be defined either at the eye angles belief or at the last projection level, each in their own domain. For example, imposing that the agent should perceive a projection of an object at the center of the camera plane results in the generation of a prediction error that ultimately moves the eyes toward that object – hence emphasizing the importance of active sensing strategies to enhance inference [32, 33].

However, the fact that there are two open pathways through which the prediction errors of the projections can flow – the eye angles and the absolute beliefs – may be problematic in some cases, e.g., during simultaneous depth estimation and target fixation. It is natural to think that depth estimation follows target fixation. In fact, top-down processing to verge on a target is generally not necessary: when an image is presented to a camera, the latter might move into this projected space directly, resulting in a simpler control [34, 35]. Then, depth could be computed directly from vergence cues. However, our model assumes that the eye angles generate the projections by first performing a roto-translation in the 2D space using the estimated depth, allowing further mechanisms for more efficient inference. Under this assumption, target fixation in the projected space is possible through a top-down process that is constantly biased by the high-level belief. Nonetheless, a direct vergence control not considered here could be implemented by additional connections between the belief over the 2D or projected points, and the angles belief.

With these considerations, it would therefore appear plausible that the two processes of depth estimation and target fixation run in parallel. But when this is the case, the prediction errors of the projections drive the two high-level beliefs independently toward a direction that minimizes the free energy, leading the agent to be stuck in an intermediate configuration with incorrect object depth and eye angles. One way to solve this issue that we pursue here consists of decomposing the task into cyclical phases of action and perception [24, 23]. During an action phase, the 2D belief is fixed and the agent can fixate on the predicted projections. Instead, during a perception phase, the agent can infer the 2D position of an object but is not allowed to move its sight. This implies that the prediction errors of the projections will alternatively flow in different directions (2D position and eye angles) one step at a time, which results in (i) the object being pulled toward the center of the camera planes and (ii) the estimated 2D position converging toward the correct one, as shown in Figure 6. Action-perception cycles have been studied in discrete time models of Active Inference; for example, cycles of saccades and visual sampling allow an agent to reduce the uncertainty over the environment (e.g., by rapidly oscillating between different points for recognizing an object) [36, 37]. Here, we show that action-perception cycles are also useful in continuous time models, such as the one used here, to ensure an effective minimization of the free energy, but also when an agent is required to reach an object with the end-effector while inferring the lengths of its limbs [23]. In summary, the two processes of recognizing a face and estimating an object’s depth can be viewed as actively accumulating sensory evidence, although at different timescales. From a brain perspective, action-perception cycles have been often associated with hippocampal theta rhythms and cortical oscillations that might segment continuous experiences into discrete elements [38, 24]. From a more technical perspective, the cyclical scheme that we propose for action-perception cycles – which consists of keeping one aspect of the optimization objective fixed when updating the other – is commonly used in various optimization algorithms such as expectation-maximization [39], and a similar approach is used for learning and inference in Predictive Coding Networks [40, 41].

Third, our results show that active vision improves depth estimation. But if vergence does not provide a useful cue for depth, how is this possible? The answer comes from the nonuniform resolution of the fovea, which has far more receptors at the center of fixation than in the peripheral vision. It is supposed that this nonlinear resolution allows gathering sensory processing resources around the most relevant sources of information [42]. Under this assumption, the best performances are achieved when both eyes fixate on the object (as shown in Figure 7). As noted in [43], when stereo cameras have a nonsymmetrical vergence angle, the error is minimum when the projections of a point fall at the center of the camera planes. Hence, vergence can effectively play a key role in depth estimation, besides providing a unified representation of the environment. This can be appreciated by considered that the *infer vergence* and *active vision* models are more accurate than the *infer parallel* model, in the nonuniform resolution condition. Rather, with a uniform resolution, the error is larger when the eyes converge to the target, because the focal angle of the pixels in the center is larger than in the periphery [43]. In addition to the increased error, the estimation seems to be further slowed down by the inference of the different reference frames due to vergence. Taken together, the consequence is that in the uniform resolution scenario the best estimation is achieved with fixed parallel eyes (see Figure 7), and active vision does not bring any advantage to the task. Since maintaining parallel eyes in the uniform condition results in a slightly higher accuracy, such simulations may be useful to understand in which cases verging on a target would improve the model performance. This might be helpful for future studies in bio-inspired robotics, especially when extending the proposed model to implement high-level mechanisms, such as: higher-level priors that result from the integration of cues from different sensory modalities; or attentional mechanisms that unify the visual sensations into a single experience.

The model presented in this study has some limitations that can be addressed in future studies. For example, we used a fixed focal length *f* during all the simulations. However, in a more realistic setting, the focal length might be considered as another DoF of the agent – and might be thus changed through suppression of proprioceptive prediction errors – speeding up the inferential process for objects at different distances. Furthermore, although the presented simulations only estimate the depth of a 2D point, it could be potentially extended to deal with 3D objects and account for vertical binocular disparity [44]. This would involve augmenting all the latent states with the new dimension and performing a sequence of two rotations as intermediate levels, before the eye projections are predicted. Then, the vergence-accommodation belief would also be extended with a new DoF, letting the agent able to fixate on 3D objects. Future studies might also investigate how the scaling factor of Equation 29 of the nonuniform resolution – but also more realistic nonuniform transformations – affect performance and help to model the human data (e.g., [25]). Also, it could be useful to adopt an off-center fovea on one of the two retinas and analyze the agent’s behavior to bring the two foveas on the target. Finally, another interesting direction for future research would be combining the architecture proposed here with a more sophisticated kinematic Active Inference model [21], for example embodying a humanoid robot with multiple DoF [45, 46, 47, 48, 49, 47]. This would allow – in contrast to state-of-the-art models that provide the agent either with the 3D environment directly as a visual observation [20], or with a latent space reconstructed from a Variational Autoencoder [50] – inference of the 3D position of an object through the projections of the eyes, which can then be used for subsequent tasks, such as reaching and grasping.

## 5 Acknowledgments

This research received funding from the European Union’s Horizon H2020-EIC-FETPROACT-2019 Programme for Research and Innovation under Grant Agreement 951910 to I.P.S., No. 945539 (Human Brain Project SGA3) to G.P. and No. 952215 (TAILOR) to G.P.; the European Research Council under the Grant Agreement No. 820213 (ThinkAhead) to G.P., the Italian Ministry for Research MIUR under Grant Agreement PRIN 2017KZNZLN to I.P.S., and the PNRR MUR projects PE0000013-FAIR and IR0000011–EBRAINS-Italy to G.P. The GEFORCE Quadro RTX6000 and Titan GPU cards used for this research were donated by the NVIDIA Corporation. The funders had no role in study design, data collection and analysis, decision to publish, or preparation of the manuscript.

